# A temperature-sensitive and interferon-silent Sendai virus vector for CRISPR-Cas9 delivery and gene editing in primary human cells

**DOI:** 10.1101/2024.05.03.592383

**Authors:** Christian S Stevens, Jillian Carmichael, Ruth Watkinson, Shreyas Kowdle, Rebecca A Reis, Kory Hamane, Jason Jang, Arnold Park, Olivier Pernet, Wannisa Khamaikawin, Patrick Hong, Patricia Thibault, Aditya Gowlikar, Dong Sung An, Benhur Lee

**Affiliations:** Department of Microbiology, Icahn School of Medicine at Mount Sinai, New York, NY 10029; UCLA School of Nursing, Los Angeles, California, 90095; UCLA AIDS Institute, Los Angeles, California, 90095

**Author notes:** Corresponding authors (BL), (DSA). Author contributions: CSS, RW, OP, DSA and BL conceived and designed the study. CSS, JC, RW, SK, RAR, JJ, AP, OP, PH, PT, AG, AB, and BK collected data. CSS, OP, JJ and KH analyzed the data. CSS wrote the original drafts of the paper. BL and DSA reviewed the draft, supported data analysis, and provided invaluable direction throughout the conceptualization and execution of the project. All authors had the opportunity to review the manuscript prior to submission.

**Keywords:** Gene editing, CRISPR/Cas9, viral vector, Paramyxoviridae, Sendai virus, hematopoietic stem and progenitor cells, HIV, CCR5

## Abstract

The transformative potential of gene editing technologies hinges on the development of safe and effective delivery methods. In this study, we developed a temperature-sensitive and interferon-silent Sendai virus (ts SeV) as a novel delivery vector for CRISPR-Cas9 and for efficient gene editing in sensitive human cell types without inducing IFN responses. ts SeV demonstrates unprecedented transduction efficiency in human CD34+ hematopoietic stem and progenitor cells (HSPCs) including transduction of the CD34+/CD38-/CD45RA-/CD90+(Thy1+)/CD49f^high^ stem cell enriched subpopulation. The frequency of *CCR5* editing exceeded 90% and bi-allelic *CCR5* editing exceeded 70% resulting in significant inhibition of HIV-1 infection in primary human CD14+ monocytes. These results demonstrate the potential of the ts SeV platform as a safe, efficient, and flexible addition to the current gene-editing tool delivery methods, which may help to further expand the possibilities in personalized medicine and the treatment of genetic disorders.

## INTRODUCTION

The bench and bedside potential of CRISPR-Cas9 gene editing is clear^1,2^, with vast implications for monogenic and infectious diseases^3–10^. In particular, a path to the clinic is likely to involve *ex vivo* editing in CD34+ hematopoietic stem and progenitor cells (HSPCs), targeting relevant diseases such as β-thalassemias^11–13^, sickle cell disease^14^, and even HIV^15^. The key to *ex vivo* gene therapies is safe and efficacious gene editing and delivery vectors. Viruses can be engineered as a vector to efficiently deliver CRISPR-Cas9 for *in vivo* and *ex vivo* editing^16–20^ due to their innate ability to enter host cells and deliver genetic material. However, the use of DNA viral vectors, such as lentivirus (LV), adeno-associated virus (AAV), and adenovirus (AdV), carries significant risks and obstacles, particularly in delivering RNA-guided endonucleases like CRISPR-Cas9.

Early use of DNA viral vectors has shown extraordinary successes^24,25^ but has also exhibited some of the serious risks associated with vector integration^26,27^ and immugoenicity^28,29^. One of the inherent obstacles in using a DNA virus is the potential for integration into the host genome^21^. For example, AAV delivery of Cas9 into mice resulted in 5% of all edited cells containing some integration of the AAV genome^22^. While the likelihood of integration can be reduced, it cannot be eliminated fully. Other, less dangerous obstacles also exist including size constraints of vectors like AAV^30^, which can be complex to circumvent^31–35^. To realize the full potential of CRISPR-Cas9 in patients, we need an efficient, safe vector that supports the viability and functionality of edited HSPCs and accommodates a flexible range of editing tools.

The use of RNA viruses, such as SeV, instead of DNA viruses as delivery vectors for CRISPR-Cas9 is one method for improving the safety of CRISPR-Cas9 editing in human cells. SeV uses sialic acid as its cellular receptor^44^ which allows it to efficiently transduce and deliver foreign genetic material to a wide variety of cell types including human CD34+ HSPCs^37^, lung airway epithelium^38^, neurons^39^, dendritic cells^40^, and many others^41–43^. SeV has other important advantages as a safe gene therapy vector: it is a non-segmented, negative-sense, RNA virus^41^ with no risk of integration into the host^45^. Additionally, it has never been linked to human disease and has been extensively studied and modified to develop temperature-sensitive (ts) and replication-defective (ΔF) vectors.

We previously published our first-generation Sendai virus vector for highly efficient Cas9-mediated editing of CCR5 (SeV-Cas9-CCR5) in human cells with minimal off-target effects^36^. In this system, guide RNA (gRNA) function depends on cleavage at the 5’ and 3’ termini of the gRNA-tracrRNA sequence. In the context of SeV, a negative sense RNA virus, we accomplished this by flanking the gRNA-tracrRNA with two ribozymes. Upon expression of the transcript, the ribozymes self-cleave and precisely liberate the gRNA. Our recombinant SeV- Cas9 virus achieved highly efficient editing in both HEK293 cells and primary human monocytes, without selection of transduced cells. These results opened the door to further develop of Sendai virus as a vector for efficient delivery of CRISPR-Cas9^36^.

Here, we report the development of a temperature-sensitive replication-restricted ts SeV- Cas9 by introducing known ts mutations in the SeV P and L genes that comprise the viral polymerase complex. These mutations potently and stably restrict SeV replication at physiological temperature and minimize interferon (IFN) responses, whilst maintaining efficient infection and replication at permissive temperatures. Notably, our ts SeV vector achieved high transduction efficiencies of in human CD34+ HPSCs, resulting in transduction of ∼90% of the CD34+/CD38-/CD45RA-/CD90+(Thy1+)/CD49f^high^ subpopulation. This population is of particular importance as single one of these cells is capable of hematopoietic reconstitution in a humanized NSG mouse^46^. Following infection, edited HSPCs maintained multilineage colony formation. We also edited primary human CD14+ monocyte-derived-macrophages with an efficiency of approximately 90% leading to *CCR5* disruption and significant inhibition of HIV infection. These results demonstrate the ts SeV platform as a promising addition to the current gene-editing tool delivery methods, which may help to further expand the possibilities of efficient gene editing in human HSPCs for the treatment of genetic disorders and infectious diseases.

## RESULTS

### Mutants in P and L genes result in a temperature-sensitive phenotype

The SeV RNA polymerase complex consists of the P and L proteins, which together are responsible for viral RNA synthesis^47^. Several mutations in both P and L are known to confer temperature sensitivity (ts), restricting viral growth to 32-34°C^48,49^ (Fig. 1A). To facilitate an improved safety profile by temperature-restricting SeV replication, we integrated these mutations into our temperature-sensitive vector (ts SeV-Cas9). We introduced three change-to-alanine mutations located in the L-binding domain of P^47^ (D433A, R434A, and K437A). In addition, this recombinant virus also has two L mutations (L1558I and K1795E) instead of just one to better restrict wild-type (wt) reversion^48^. These mutations confer a ts phenotype while maintaining sufficient vector titers^48,50^.

**Figure 1.**
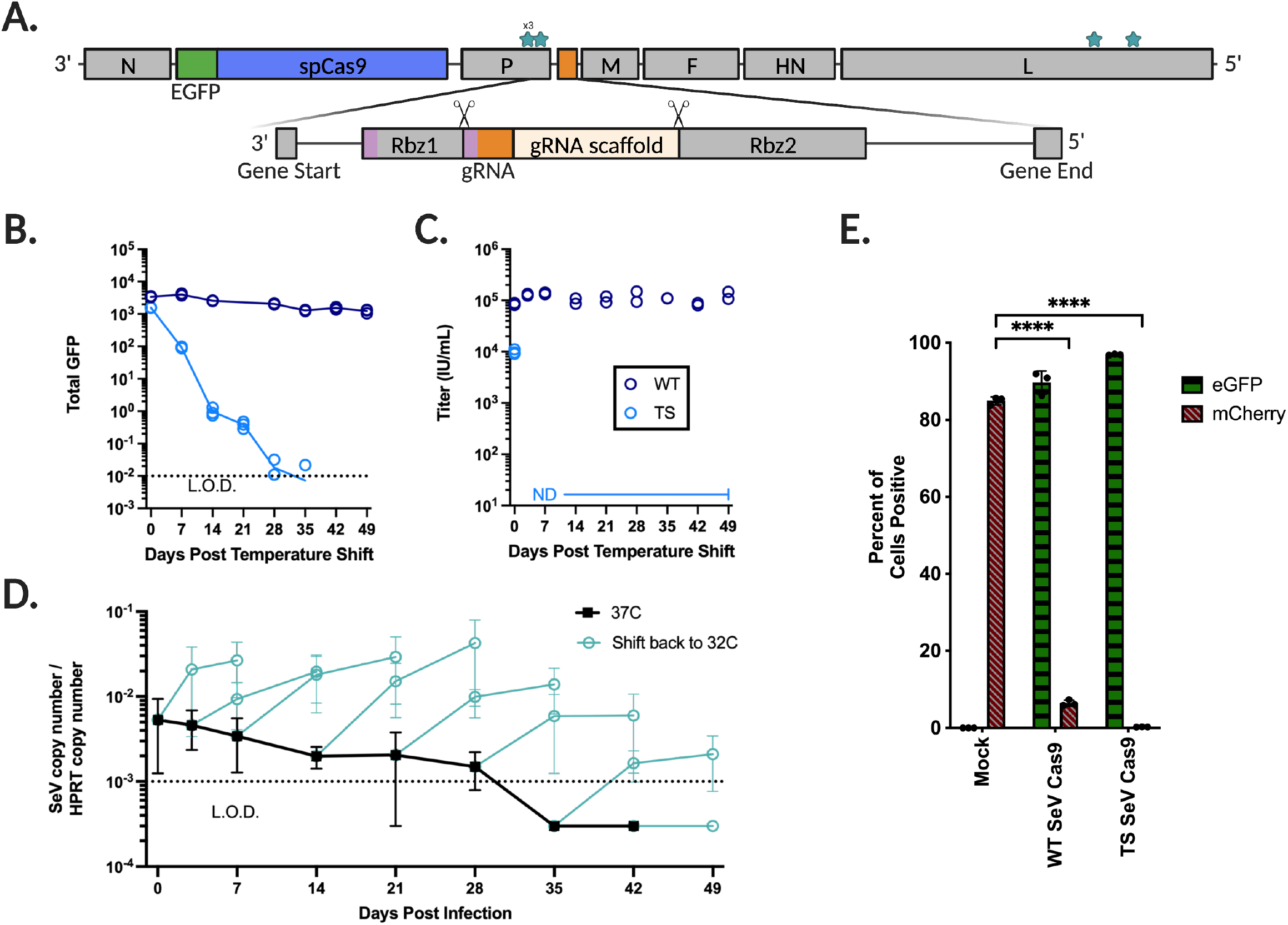
Sendai virus incorporating Cas9 and a guide RNA flanked by self-cleaving ribozymes contains mutations in P and L that impart a temperature-sensitive phenotype. **(A)** Shown is the Sendai virus genome containing SeV genes N (nucleoprotein), P (phosphoprotein), M (matrix), F (fusion protein), HN (attachment protein), and L (large RNA- dependent RNA polymerase). An eGFP-P2A-Cas9 cassette (5.1 kb) was inserted between N and P, and a guide RNA flanked by self-cleaving ribozymes (rbz 1 and 2) (0.2 kb total) was inserted between P and M (see Materials and Methods for further details). Mutations were made in both P and L in order to impart a temperature-sensitive phenotype. **(B)** 293T cells are infected at 32°C for two days by either wild type (WT) or temperature sensitive (TS) SeV-Cas9 then shifted to 37°C. Shown is the total GFP measured over 7 weeks, performed in triplicate. The dotted line indicates the limit of detection as determined by the mean of mock-infected cells. **(C)** As in B, supernatant is taken and used to infect Vero-CCL81 cells and the titer is calculated in infectious units per mL. Experiment performed in duplicate. **(D)** 293T cells are infected at 32°C for two days then shifted to 37°C (black) until the timepoint indicated (cyan). Experiment performed in technical triplicate and biological duplicate. **(E)** Both wt and ts SeV-Cas9 containing a gRNA targeting mCherry are used to infect 293 FLP-mCherry cells. eGFP positivity indicates successful transduction of the SeV-Cas9 and mCherry indicates a lack of indels and subsequent knockout of the mCherry gene. Performed in triplicate. Editing compared using Welch’s t test. (ns, not significant; **, p < 0.01; ***, p < 0.005, and ****, p < 0.0001).

We infected 293T cells at 32°C and incubated them for 48 hours before shifting the temperature to 37°C. Following the temperature shift, SeV gene expression rapidly declined and eventually became undetectable (Fig. 1B). Moreover, there was no detectable virus titer for ts SeV-Cas9 seven days after the temperature shift (Fig. 1C). Notably, ts SeV-Cas9 viral genomes remained detectable by RT-qPCR in some samples until five weeks at 37°C, but by the sixth week, the virus was unable to recover (Fig. 1D). Crucially, at 32°C, ts SeV-Cas9 demonstrated equal or superior efficiency in entering 293T cells stably expressing mCherry (293T-mCh) and subsequently editing mCherry compared to wt-SeV-Cas9 (Fig. 1E). Although editing efficiency was relatively low after only two days at 32°C, the percentage of edited cells continued to rise for five days following the shift to 37°C (Supp. Fig. 1).

We are therefore able to show that our ts SeV-Cas9 vector, with its mutations in the P and L genes, displays a temperature-sensitive phenotype that allow for efficient editing and clearance following the editing process. This temperature-sensitive vector provides a potential approach to minimizing cytotoxic effects associated with sustained viral infection while maintaining the ability to efficiently edit target genes.

### Infection by ts SeV-Cas9 elicits an IFN-*silent* phenotype

Given the desire for clearance and a minimal impact on infected cells apart from gene editing, we next interrogated whether ts mutation can minimize interferon responses. While examining the persistent effects of ts SeV infection in cells after shifting to 37°C, we assessed the transcriptional upregulation of two interferon-stimulated genes (ISGs): RIG-I and IFIT1 following infection of 293T cells. We discovered that ts SeV infection resulted in significantly lower expression of both RIG-I and IFIT1 in infected cells, compared to those infected by wild-type (wt) SeV (Fig. 2A,B). Cells were infected with either ts or wt SeV at 34°C, and after two days, the temperature was shifted to 37°C. The difference in ISG expression was most prominent immediately after infection but persisted for at least 30 days following the shift to 37°C.

**Figure 2.**
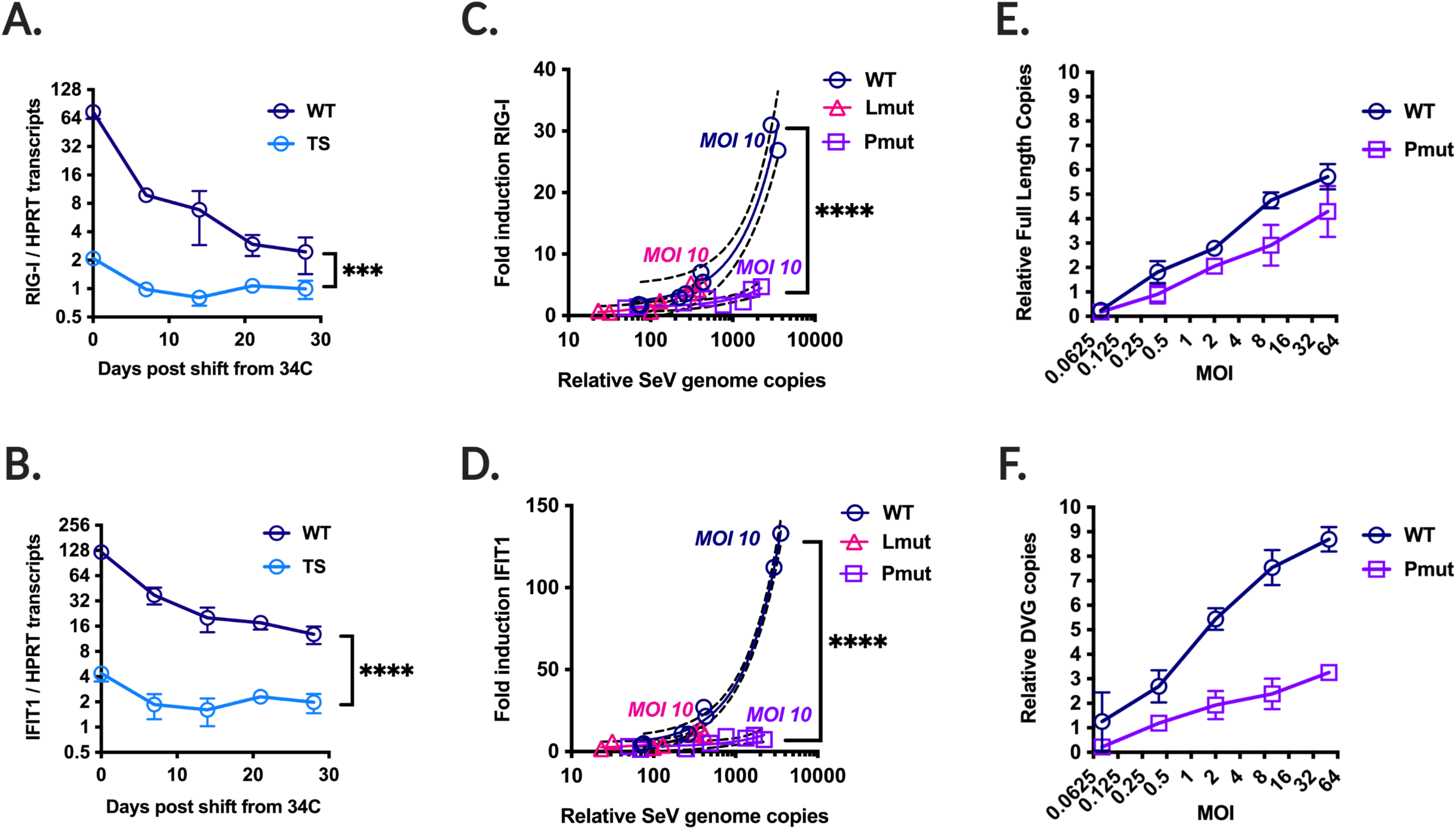
The impact of mutations in P and L on Sendai virus ISG stimulation. Sendai virus was used to infect 293T cells. **(A)** and **(B)** wt and ts SeV are infected at 34C for two days then shifted to 37C. (A) RIG-1 transcripts or (B) IFIT1 transcripts are normalized by HPRT transcripts as determined y RT-qPCR are measured. **(C)** and **(D)** Sendai virus containing the temperature-sensitive mutations in P only, L only, or wild type is used to infect 293T cells at multiple MOIs. Each MOI is performed in triplicate and relative SeV genomes are compared against the fold induction of (C) RIG-I or (D) IFIT1. **(E)** and **(F)** Both WT and Pmut SeV are used to infect 293Ts are 5 different MOIs and both (E) relative full length copies and (F) relative DVG copies are measured. All experiments performed in technical triplicate, error bars indicate SEM, and all comparisons done using Welch’s t test. (ns, not significant; **, p < 0.01; ***, p < 0.005, and ****, p < 0.0001).

To isolate the mutations responsible for the differential expression of ISGs, we created SeV variants containing just the mutations in P (Pmut; D433A, R434A, and K437A) or just the mutations in L (Lmut; L1558I and K1795E). We infected 293T cells with these SeV variants at five different multiplicities of infection (MOIs) and evaluated RIG-I and IFIT1 fold induction after two days at 34°C to determine if the reduction in ISG expression could simply be attributed to the P and/or L mutations attenuating viral growth. We found that Lmut was highly attenuated, with fewer SeV genome copies at the same MOIs compared to both wt and Pmut, but Lmut induced ISG expression similarly to wt SeV at comparable genome copy levels (Fig. 2C,D).

However, there was no significant difference in the relative number of SeV genome copies at a given MOI between wt SeV and Pmut SeV. Despite this, we observed a significant increase in the fold induction of both RIG-I and IFIT1 following infection by wt SeV compared to Pmut SeV (Fig. 2C,D). This suggests that Pmut SeV exhibits an interferon-silent phenotype that cannot be explained simply by attenuation as that would be reflected in a difference in the SeV genome copy number at the same MOI.

To further investigate the differences between wt SeV and Pmut SeV, we measured both full-length SeV genome copies and defective viral genomes (DVGs) using RT-qPCR for copy-back genomes. DVGs are known to be immunostimulatory^51^, therefore a reduction in DVG production might contribute to the observed IFN-silent phenotype. Our data revealed no significant difference in relative full-length SeV genome copies as measured by area under the curve (Fig. 2E). However, when examining relative DVG copies, we found significantly fewer relative copies in Pmut SeV compared to wt SeV (Fig. 2F). This suggests a potential role for DVG production in ISG stimulation during SeV infection. Furthermore, specific residues in P which confer temperature sensitivity also contribute to DVG production and mutation of these residues decreases DVG production resulting in an interferon-silent phenotype.

### ts SeV-Cas9 can effectively deliver either one or two guides in a single construct

Our 1^st^ generation SeV-Cas9 expressed a guide RNA and Cas9 via the creation of two additional cassettes^36^. The first cassette was inserted between N and P and includes eGFP- P2A-Cas9 (5.1 kb). The other cassette is between P and M and contains a guide RNA flanked by self-cleaving ribozymes (0.2 kb) (Fig. 1A and 3A). Efficient editing requires the cleavage of the gRNA and scaffold from the capped and polyadenylated viral mRNA^36^. For ribozyme self-cleavage to occur, the ribozyme must fold consistently. As shown in Fig. 3A, cleavage of the gRNA occurs upstream of one half of Rbz1’s stem-loop. This means that the first six bases in the chosen gRNA must be a perfect reverse compliment to the most upstream six bases of Rbz1. Therefore, for each new guide incorporated into our SeV-Cas9 system, we must also uniquely adjust Rbz1.

**Figure 3.**
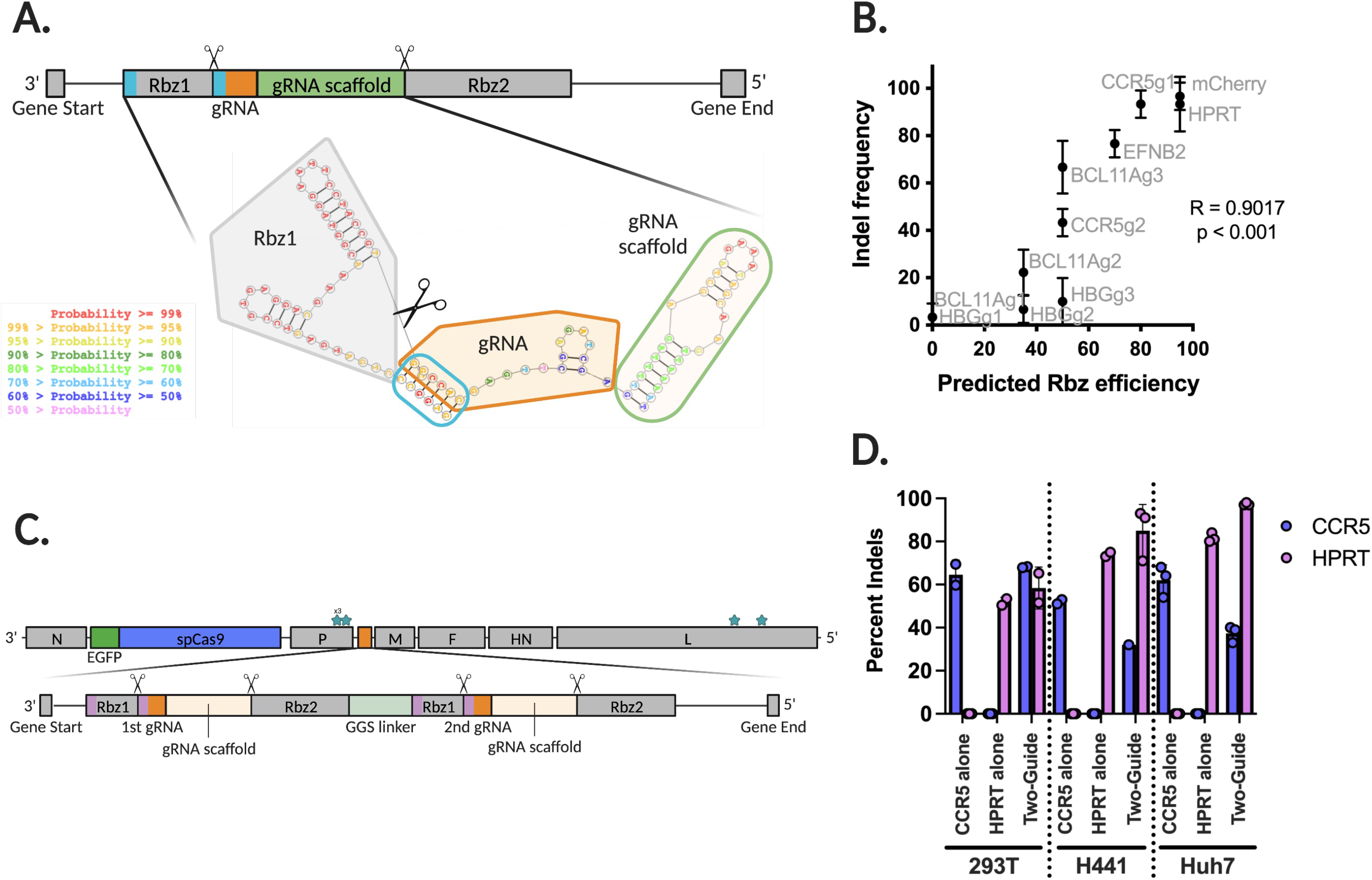
SeV-Cas9 can deliver a diversity of guides and can utilize novel guide strategies. (A) The gRNA cassette in SeV-Cas9 contains two ribozymes, both necessary for efficient downstream editing. Using RNAstructure we visualize and calculate the probability of proper stem-loop formation required for ribozyme cleaved. **(B)** We compare the indel frequency measured by Sanger sequencing against the predicted ribozyme efficiency as calculated using information from RNAstructure. Significance shown that slope does not equal zero. **(C)** The gRNA cassette in SeV-Cas9 capable of delivering two separate gRNAs by flanking both with two ribozymes each, separated by a GGS linker. **(D)** Comparing the single guide systems targeting CCR5 or HPRT and the two-guide system targeting both in 293Ts, H441s, and Huh7 cells. Experiment performed in triplicate and indels calculated via Sanger sequencing and Synthego ICE analysis.

We rescued 11 different ts SeV-Cas9 viruses containing a single gRNA targeting one of five different genes (*HBG*, *BCL11A*, *CCR5*, *EFNB2*, and *mCherry*). We found that these previously tested and otherwise efficient guides displayed highly variable editing efficiency in the context of our SeV-Cas9 system (Fig. 3B). Because editing efficiency might be affected by gRNA cleavage efficiency via our self-cleaving ribozymes, we tested whether predicted ribozyme folding efficiency would correlate with editing efficiency. Using the RNAstructure^52^ RNA-folding prediction webserver, we extracted the most likely RNA structure for each gRNA cassette (Fig. 3A). Within that structure we looked for the existence of predicted base pairing at the expected stem-loop. We then took the mean of the predicted probability of basepair formation for each of the six pairs (assigning a zero in the case of no predicted pairing) in order to calculate a value we called “predicted Rbz efficiency.” Using indel frequency as a proxy for gRNA editing efficiency we observed a positive relationship between gene editing frequency and predicted Rbz efficiency (Fig. 3B).

To add to our current toolset, we expanded the gRNA cassette to include two different gRNAs instead of one. This was accomplished using two sets of flanking self-cleaving ribozymes in a single cassette (Fig. 3C). To test the efficiency of this system, we incorporated two different guides, one against CCR5 and the other against HPRT, and compared editing across three different cell lines. We then measured the occurrence of editing within both *CCR5* and *HPRT* in cells infected with ts SeV-Cas9 CCR5 (CCR5 alone), ts SeV-Cas9 HPRT (HPRT alone), and ts SeV-Cas9 CCR5/HPRT (two-guide) (Fig. 3D). We observed editing of both targets, either individual or in combination, in our one and two-guide systems respectively.

### Efficient ts SeV-Cas9-CCR5 mediated transduction in CD34+ hematopoietic stem progenitor cells (HSPCs)

Gene editing in human HSPCs has significant scientific and clinical potential for treating many diseases. In particular, efficient *CCR5* editing in human HSPC has a great potential for developing HIV therapies. We therefore investigated ts SeV-Cas9-mediated *CCR5* editing in CD34+ HSPC. CD34+ cells derived from human fetal liver and G-CSF mobilized peripheral blood. HSPC derived from multiple donors (n=3) were transduced with ts SeV-Cas9-CCR5 at various multiplicity of infections (MOI = 0.1 to 20) for one to 20 hours. Transduced cells were cultured at 34°C for 3 days and analyzed for %eGFP expression in CD34+ HSPCs by flow cytometry. ts SeV-Cas9-CCR5 transduced human fetal liver (FL) derived and mobilized peripheral blood (mPB) CD34+ HSPC consistently at >90% eGFP+ within 3 days at MOIs greater than one at 34°C (Fig. 4A,C,D). The length of incubation with virus (1hr or 20hrs) prior to changing the media had no effect on transduction efficiency. Moreover, we observed transduction efficiency of >95% in CD34+/CD38-/CD90+(Thy1+)/CD49f^high^ hematopoietic stem cells (HSCs) (Fig.4B), the HSPC sub-population capable of hematopoietic reconstitution by a single cell in a hu-NSG mouse^53^. These results demonstrate unprecedented transduction efficiency of ts SeV-Cas9-CCR5 in CD34+ HSPC.

**Figure 4.**
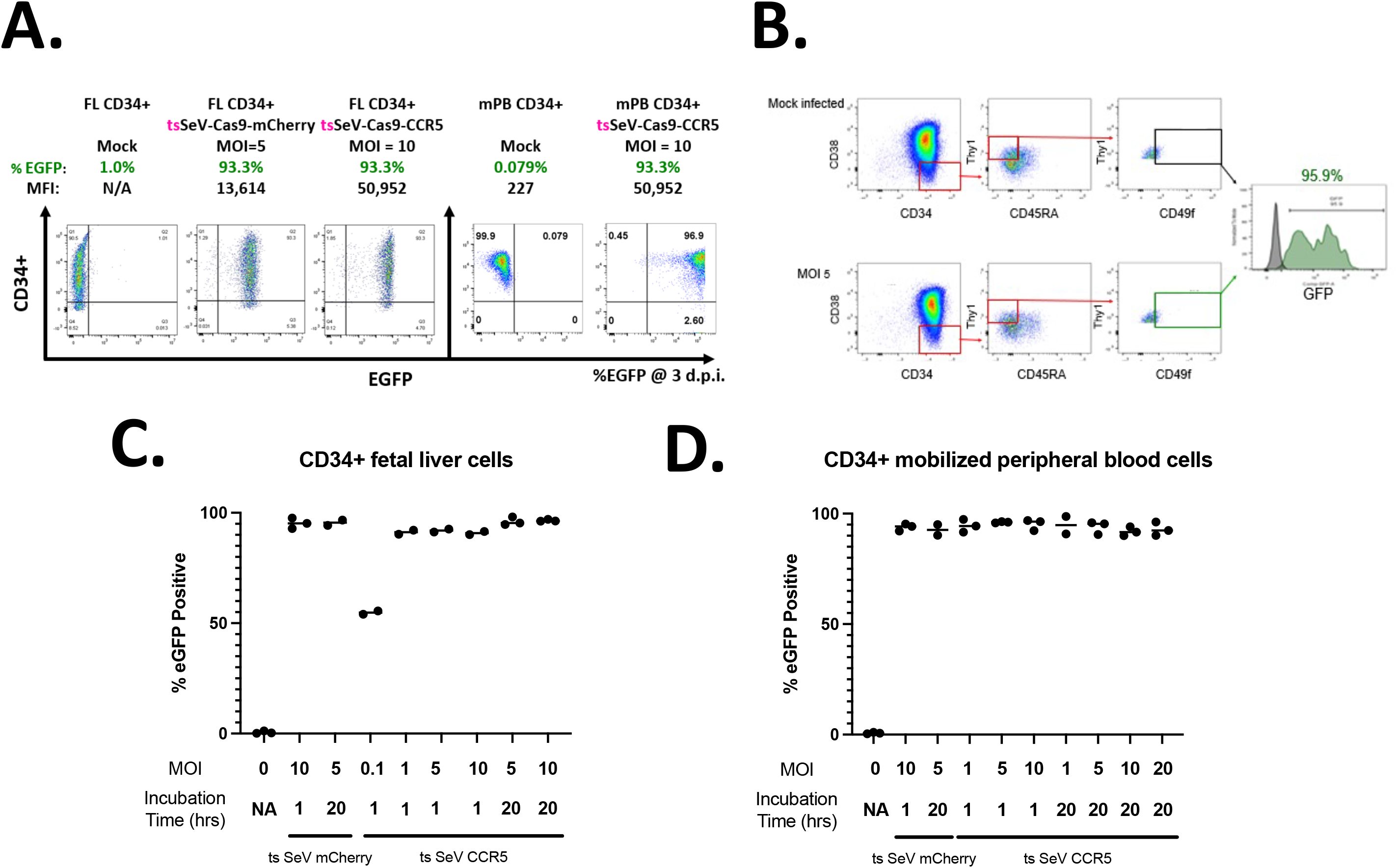
Efficient CD34+ HSPC transduction by the ts rSeV-Cas9-CCR5. (A) Representative flow cytometry data of human fetal liver and G-CSF peripheral blood mobilized CD34+ HSPC infected with ts rSeV-Cas9-CCR5 at MOI 5 and 10 at 34°C. Flow cytometry showed >90% transduction (EGFP+) relative to mock infected cells at 3 dpi. **(B)** Efficient transduction in the rare human CD34+/CD38-/Thy1+/CD49f ^high^ HSC enriched subpopulation. Percent eGFP (95.9%, green histogram) was determined relative to mock (gray histogram) infected cells. **(C)** and **(D)** CD34+ HSPC transduction by ts rSeV-Cas9-CCR5 yielded >90% transduction across all MOIs greater than 1 tested, both in (C) fetal liver (FL) CD34+ HSPCs and (D) mobilized peripheral blood (mPB) CD34+ HSPCs.

### On-target editing efficiency of ts rSeV-Cas9-CCR5 and its effects in CD34+ HSPC

We assessed the editing efficiency of ts SeV-Cas9-CCR5 in both FL and mPB CD34+ HSPCs and show that the rate of insertion and deletion introduction within the target site *CCR5* (% indel) rises with increasing MOI. Transduction efficiency for both ts SeV-Cas9-mCherry and ts SeV-Cas9-CCR5 were >90%, but there was no significant editing of *CCR5* detected when infecting with an MOI < 1 or when a guide against *CCR5* was not present (Fig. 5A). We see efficient editing in CD34+ HSPCs starting at an MOI of 10, and therefore we used this MOI for all following experiments.

**Figure 5.**
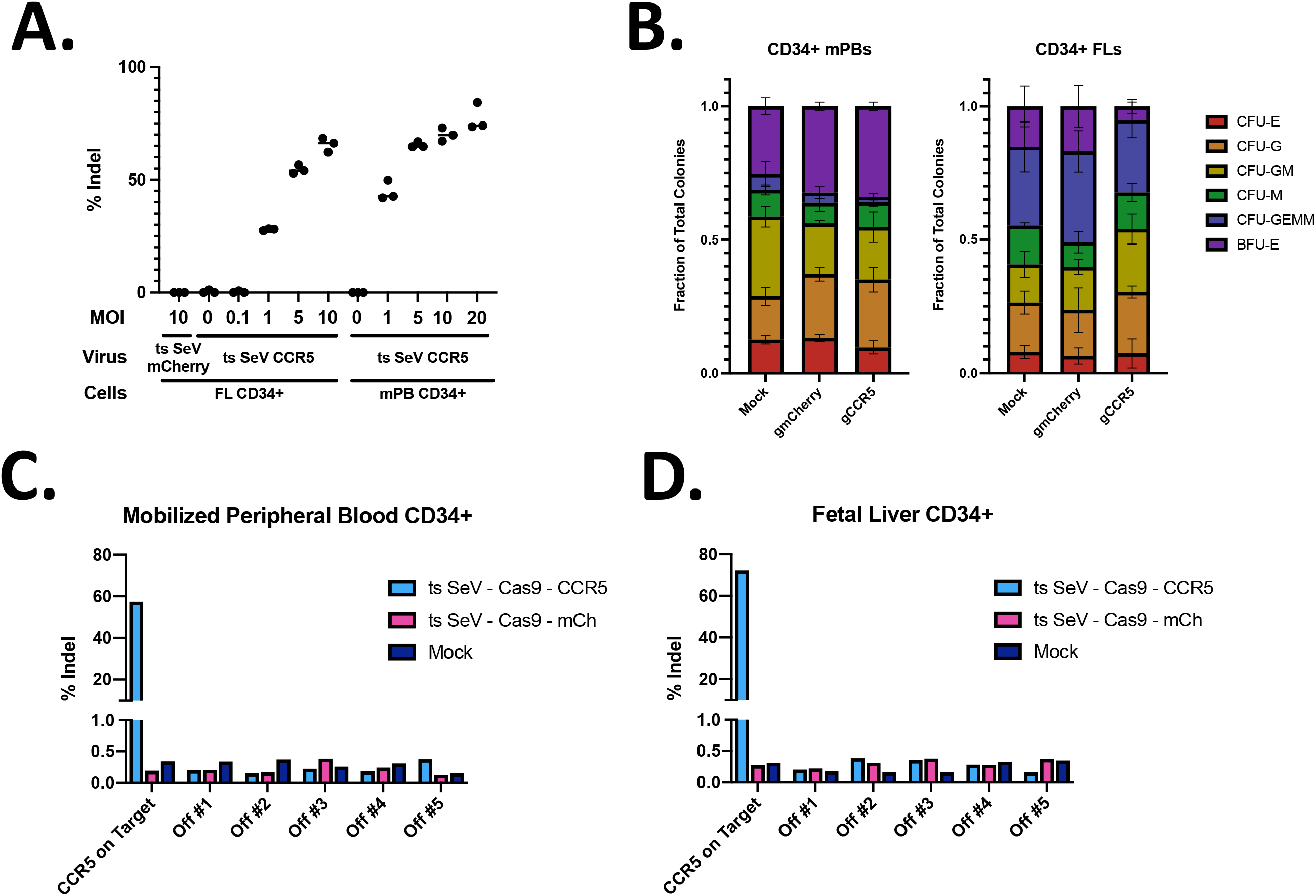
Editing efficiency in CD34+ HSPCs and the effect on hematopoietic differentiation. (A) Fetal liver (FL) CD34+ HSPCs or mobilized peripheral blood (mPB) CD34+ HSPCs were infected at multiple MOIs, with percent indels calculated via Synthego ICE analysis. **(B)** Downstream colony differentiation after ts SeV-Cas9 transduced mPB or FL CD34+ HSPCs at an MOI of 10. We measure CFU-E: CFU erythroid; CFU-G; CFU granulocytes; CFU-GM: CFU granulocytes and macrophages; CFU-GEMM: CFU granulocyte, erythrocyte, monocyte, megakaryocyte; BFU-E: Burst-forming unit-erythroid, CFU-M: Colony Forming Unit - monocytes. Error bars are SD. For raw counts see Supp. Fig. 2. **(C)** Mobilized peripheral blood CD34+ HSPCs and **(D)** fetal liver CD34+ HSPCs are infected by ts SeV Cas9 containing a guide targeting CCR5 or mCherry at an MOI of 10. Percent indels calculated via Illumina sequencing (see Materials and Methods). Top 5 off-target sites predicted by the CRISPR design tool (crispr.mit.edu). For raw counts see Supp. Fig. 2.

While we see efficient transduction and editing of CD34+ HSPCs from donors, it is vital these edited cells still hold potential for multi-lineage hematopoietic differentiation. Following mock infection as well as infection by both ts SeV-Cas9 targeting mCherry and ts SeV-Cas9 targeting *CCR5* we see similar ratios of colony formation within a single donor between CFU-E, CFU-G, CFU-GM, CFU-GEMM, BFU-E, and CFU-M although there are some differences seen between the two donors (Fig. 5B). As expected, we do see some reduction in the total number of hematopoietic colonies in the ts SeV-Cas9-mCherry relative to the mock infection, and a further reduction for ts SeV-Cas9-CCR5 (Supp Fig. 2). Presumably, the decrease in colonies in ts SeV- Cas9-mCherry transduced HSPCs is due to the effect of transduction alone as the delivered gRNA has no target in these cells. The additional colony decrease after transduction with ts SeV-Cas9-CCR5 likely results from the additional burden of double-stranded breaks. We are therefore able to show efficient transduction, editing, and downstream differentiation in CD34+ HSPCs.

We also assessed the frequency of editing at predicted off-target sites in FL and mPB derived CD34+ HSPC after transduction with ts SeV-Cas9-CCR5. Vector transduction efficiency was >90% and CCR5 editing was >88%. The frequency of editing at 5 predicted off-target sites was determined by deep sequencing and found to be <0.4% in FL-derived CD34+ HSPC and <1% in mPB-derived CD34+ HSPC (Fig. 5C,D). The occurrence of editing at these predicted off-target sites did not exceed the frequency of editing at the same sites after transduction with rSeV- Cas9-mCherry (<0.2%). These results demonstrate that ts SeVCas9-CCR5 mediates editing of CCR5 in CD34+ HSPC with minimal off-target effects.

### Efficient CCR5 editing of primary human CD14+ monocytes by ts SeV-Cas9 inhibits HIV infection

Having demonstrated the efficacy of ts SeV in delivering Cas9 and a gRNA to CD34+ HSPCs, we extended our investigation to its potential in the context of primary CD14+ monocytes for the purpose of inhibiting HIV infection. Primary CD14+ monocytes were isolated from donor blood and subsequently infected with ts SeV-Cas9-US11 or ts SeV-Cas9-CCR5 at a MOI of 10. Our guide targeting US11, a human cytomegalovirus gene, was chosen as a negative control because it has no significant off-target editing sites predicted in the human genome nor the HIV viral genome. The cells were incubated at 34°C for two days, then the monocyte-derived macrophages (MDMs) were shifted to 37°C. CCR5 editing efficiency was measured at 10 and 18 days post infection (dpi). This analysis demonstrated an editing efficiency at the *CCR5* gene of approximately 80% at 10 dpi and nearly 90% by 18 dpi in ts SeV-Cas9-CCR5 transduced cells. As expected, no *CCR5* editing was observed in the control group infected with a virus targeting US11 (Fig. 6A).

**Figure 6.**
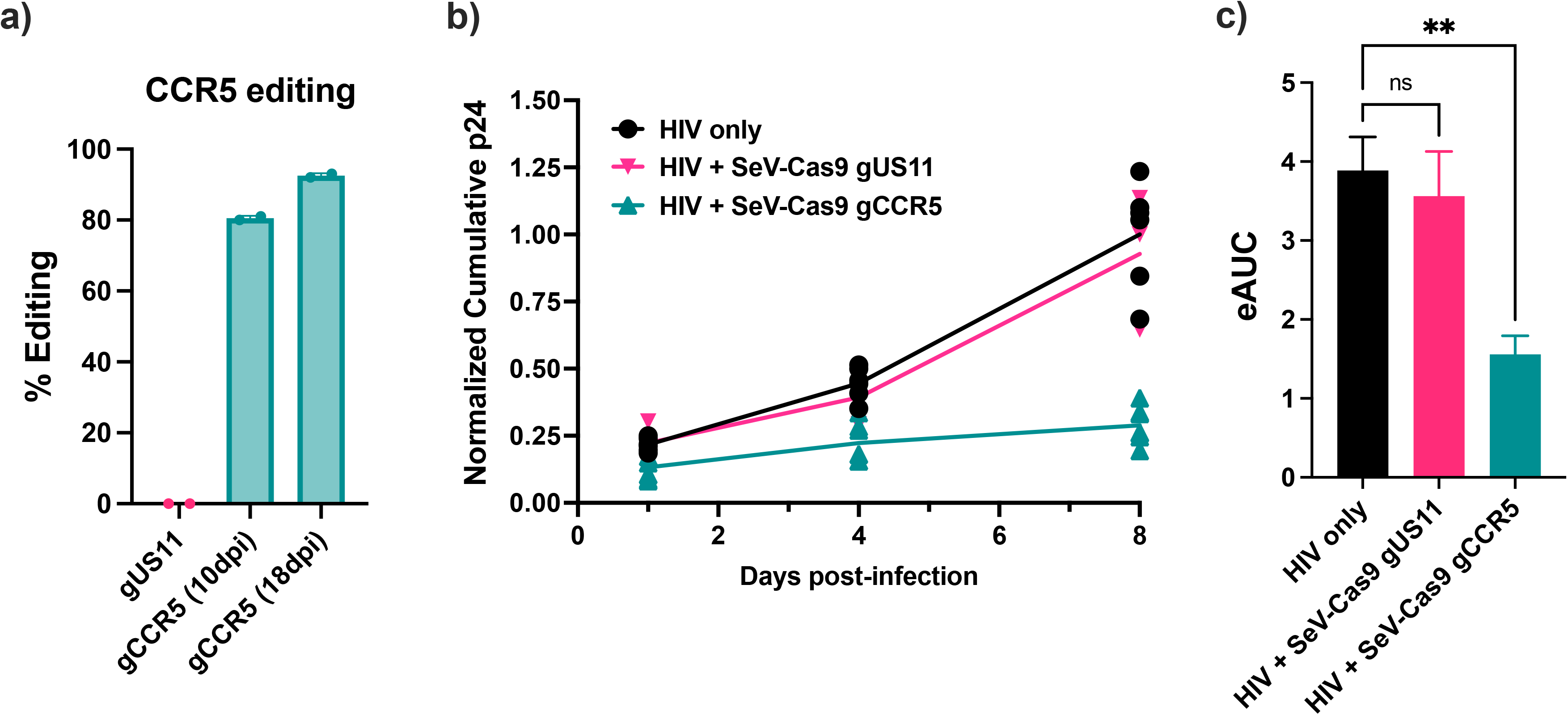
CCR5 editing of primary CD14+ monocytes with ts-SeV-Cas9 limits infection with HIV. **(A)** ts SeV-Cas9 mediated editing efficiency of CCR5 in CD14+ monocytes was determined at 10 and 18 dpi with ts-SeV-Cas9. Cells infected with a ts-SeV-Cas9 mCherry targeting virus were used as a negative control. 2 samples per condition were measured **(B)** An HIV growth curve of ts-SeV-Cas9 infected MDMs measuring the accumulation of P24 in the supernatant. Samples were collected at 1, 4, and 8 dpi and the cumulative level of P24 in each sample was calculated. Samples are from 6 replicates from HIV only, 3 replicates HIV + SeV- Cas9 US11, and 7 replicates from SeV-Cas9 CCR5. p24 values were normalized as a fraction of the average Day 8 cumulative p24 value across all replicates in a batch. **(C)** The estimated area under the curve (eAUC) was calculated for the experiment shown in B). Statistical significance was determined with Brown-Forsythe and Welch ANOVA tests.

To study the effectiveness of ts SeV-delivered CRISPR-Cas9 as a tool for inhibiting HIV infection in primary MDMs via targeting of *CCR5*, MDMs were transduced with ts SeV-Cas9- CCR5 and subsequently infected with the R5-tropic HIV strain JR-FL. Over the course of 8 days, cell supernatant was collected and analyzed for P24 to determine HIV infection (Fig. 6B). At 8 dpi there was a marked reduction in P24 accumulation in MDMs transduced with ts SeV- Cas9-CCR5 compared to cells transduced with ts SeV-Cas9-US11 or cells infected with HIV alone.The estimated area under the curve (eAUC) was calculated for each condition to determine the significance of these findings (Fig. 6C). The MDMs infected with ts SeV-Cas9- CCR5 before being challenged with HIV showed a significant reduction in P24 relative to the HIV only group (p < 0.01) but the MDMs infected with ts SeV-Cas9-US11 did not. These results indicate that the use of ts SeV-Cas9 for *CCR5* editing in primary CD14+ monocytes can lead to a reduction in HIV infection, as measured by P24 accumulation.

## DISCUSSION

The ongoing development and refinement of delivery systems for gene editing tools represent a key effort in realizing their potential for gene therapy and personalized medicine. In this study, we focused on the engineering of a temperature-sensitive Sendai virus (ts SeV) to effectively deliver the CRISPR-Cas9 system into a variety of cell types, with a particular emphasis on primary human CD34+ hematopoietic stem and progenitor cells and primary human CD14+ monocyte-derived-macrophages. Intriguingly, our modifications to the virus, intended to impart a temperature-sensitive phenotype, resulted in an additional, unexpected phenotype—interferon- silence—which could have significant implications for the safety profile and efficiency of gene editing modalities.

While SeV has been used as a clinical tool for decades^41^, it typically has several limitations. The first is that SeV has the potential to be highly immunogenic and can even induce apoptosis^45,54^. It has also been shown to induce significant IFN-γ production^55^ which can result in CD34+ HSPC depletion, impaired proliferation, and impaired self-renewal^56–59^. This is particularly important to be aware of because toxicities during gene editing delivery have been shown to result in poor engraftment in NSG mice (4%)^60^. It was the knowledge of these potential shortcomings of the system that pushed us to investigate the interferon stimulation following infection. However, the interferon-silent phenotype we observed was not expected.

In this manuscript we showed that the engineered mutations in the P and L proteins of ts SeV confer both temperature sensitivity and an interferon-silent phenotype. Specifically, we isolate the P mutations as driving the interferon-silent phenotype and the L mutations as predominantly attenuating viral growth. Here, we propose a potential explanation for the dual temperature-sensitive and interferon-silent effect by suggesting that lower protein folding constraints at 32- 34°C might increase phenotypic space. This in turn might allow the mutated P protein to do one of two things, 1) enhance viral RNA polymerase processivity or 2) otherwise antagonize host innate immune sensing and signaling pathways. If the P mutations primarily result in enhanced polymerase processivity, a potential consequence could be a decrease in the production of DVGs as shown, which are known for their immunostimulatory effects. This would explain the interferon-silent phenotype observed, although we acknowledge that further experimental validation is necessary to substantiate these hypotheses.

Through our ts SeV-Cas9 system, we demonstrated the delivery of a range of guide RNAs, highlighting the potential for expanding the applications of CRISPR-Cas9 gene editing. This system’s versatility was further demonstrated by its ability to infect and edit multiple cell types, using both single and dual-guide systems, showcasing the ts SeV-Cas9 system’s potential for flexibility and adaptability in gene editing. We also showed that our system is flexible enough to target nearly any gene because it can accommodate gRNAs previously shown to have efficient editing by computationally predicting which ribozymes pair best with a given guide. Developing a predictable and reliable system was a non-trivial step considering the inherent difficulties in delivering precisely cleaved gRNAs using an RNA virus.

Sendai virus as a vector shows significant potential, in part because of some of the constraints on other successful viral vector. The first major obstacle to other viral vectors is efficient packaging of the editing modality when size is a factor. Adeno-associated viruses (AAVs) are among the most popular viral vectors, but due to their size, ∼20 nm, they can package at most 4.5-5.2 kb of additional genetic material^30^. Packaging SpCas9 and a sgRNA requires approximately 4.2 kb of space leaving little room for any additional material. Another potential obstacle is immunogenicity which has been seen in vectors such as adenoviruses and lentiviruses^18,61,62^. In addition, because viral vectors tend to be DNA viruses, there exists a risk of integration into the host genome^21,63^. This is especially true when delivering editing tools that result in double-stranded breaks (DSBs) leading to non-homologous end joining (NHEJ) or homology-directed repair (HDR). *In vivo* in mice and macaques, AAV vector genome integration was found at the targeted site for a DSB in greater than 5% of all cells edited^22,23^. Each vector has a unique set of advantages and disadvantages, but there remains a need for an efficient and flexible vector with no risk of integration into the host genome like Sendai virus.

Perhaps one of the most important results shown here was the successful transduction of CD34+ HSPCs with the ts SeV-Cas9 system. We showed that we can leverage the clearance potential of our temperature-sensitive vector as well as its interferon-silent phenotype, to infect highly sensitive CD34+ HSPCs. We achieved transduction efficiencies of ∼90% in fetal liver derived and peripheral blood mobilized CD34+ HPSCs as well as the CD34+/CD38-/CD45RA-

/CD90+(Thy1+)/CD49f^high^ subpopulation^46^. The system showed promising transduction efficiencies, and importantly, the edited HSPCs retained their multilineage colony formation potential, a critical factor for any potential clinical applications. However, we are mindful of the considerable work still required to translate these initial findings. Given the need to continue to maximize survival, proliferation, and self-renewal in order to optimize engraftment, we recognize that there is space for additional mitigation of the cytotoxic effects inherent to transduction and NHEJ repair of double stranded breaks.

An important use case for our ts SeV-Cas9 system is infection and editing of primary human CD14+ monocytes, with the aim of inhibiting HIV infection. We employed our system to deliver Cas9 and a gRNA targeting CCR5 into these monocytes and subsequently differentiated them into MDMs. Our results were highly promising. We observed a significant editing of CCR5, reaching approximately 80% at 10 dpi, and nearly 90% by 18 dpi. To ensure the expected functional implications of CCR5 editing, we challenged the MDMs with the R5-tropic HIV strain JR-FL and monitored HIV infection over an 8-day period. In the MDMs infected by ts SeV-Cas9 CCR5, that had undergone significant CCR5 editing, we observed a marked reduction in P24 accumulation compared to the control group, demonstrating a significant inhibition of HIV infection. These findings underscore the potential of the ts SeV-Cas9 system both in the context of HIV gene editing approaches and also simply as an effective delivery tool for gene knockout with functional implications.

The ts SeV system shows significant potential as a delivery vehicle for gene editing modalities. When delivering Cas9, it is capable of transducing and editing highly sensitive cell types with functional implications. Given the pleomorphic nature of Sendai virus, efficient rescue and viral growth might be possible even when packaging increasingly large transgenes. This could potentially allow for the incorporation of other editing modalities, such as adenine base editors and prime editors, extending the utility of the system. It would also improve the safety profile of the system, further reducing cell cytotoxicity if modalities were packaged that did not induce double stranded breaks.

In summary, this study contributes to the ongoing endeavor to develop safer and more efficient delivery methods for gene editing in sensitive cell types. While the results are encouraging, they represent a step in the broader, complex landscape of *ex vivo* gene editing research. We believe that the potential of the ts SeV-Cas9 system, its future refinement, and its implications for gene therapy and personalized medicine warrant continued exploration and investigation.

## METHODS

### Maintenance and generation of cell lines

Flp-In T-REx HEK293 cells (Invitrogen, Waltham, MA), Vero cells (ATCC CCL-81), Huh7 cells (JCRB Cell Bank, JCRB0403), BSR-T7 cells (BHK- based cell line with stable expression of T7 polymerase)^64^ were maintained in Dulbecco’s modified Eagle’s medium (Invitrogen) supplemented with 10% heat-inactivated fetal bovine serum (FBS) (Atlanta Biologicals, Flowery Branch, GA). Flp-In T-REx HEK293 cells were additionally maintained in blasticidin and ZEOCIN according to manufacturer protocol. mCherry-inducible cells were generated as previously described^36^. In brief, the mCherry gene was cloned into pcDNA5/FRT/TO and transfected with pOG44 containing Flp-recombinase into parental Flp-In T-REx HEK293 cells. Cells were then put under selection with hygromycin and blasticidin according to manufacturer resulting in doxycycline-inducible expression of mCherry. H441 (NCI- H441 ATCC) cells were maintained in Roswell Park Memorial Institute (RPMI) 1640 medium (Invitrogen) supplemented with 10% heat-inactivated FBS (Atlanta Biologicals).

### Cell surface expression and receptor binding by flow cytometry

Cell surface expression of RBP was assessed by transfecting the respective wild type or mutant constructs into HEK293T cells with BioT (Bioland Cat. No. B01-01). Two days post transfection, the cells were gently collected with 10mM EDTA to avoid cleavage of the glycoprotein. Cells were then stained with a 1:2000 dilution of anti-HA (Gentex Cat. No.). For assessing receptor binding, human Fc (hFc) tagged soluble EFNB2 (sEFNB2-hFc) from R&D (Cat. No. 7397-EB-050) or sEFNB3-hFc (Cat. No. 7924-EB-050) was used. Cell surface expression of EFNBs on newly generated cell lines was verified by seeding cells in a 12 well plate prior to collecting with 10nM EDTA and staining with sEPHB3-hFc (R&D Cat. No 5667-B3-050). The Attune was used for all flow cytometry data acquisition and data were analyzed using FlowJo software.

### Assaying defective viral genome production

Virus was produced by first infecting BHK-21 cells, lacking functional type I IFN. We then titer supernatants on IFN-competent HEK293T cells. We then RT-qPCR for full-length SeV genomes and DVGs (assay adapted from Xu 2017).

### Sendai virus construction and rescue

Design, construction, and rescue of rSeV-Cas9 was performed as previously described^36^. In brief, we used a RGV0 (kind gift of Nancy McQueen) derived Fushimi strain Sendai virus with an eGFP reporter between N and P via duplication of the N to P intergenic region^65,66^ and mutations in F and M allowing for trypsin-independent growth^67^. *S. pyogenes* Cas9 was amplified from px330 (Addgene, cat #42230, Feng Zhang) and linked with a P2A ribosomal skipping sequence to eGFP in rSeV. The gRNA cassette was inserted between the P and M genes via duplication of the P-to-M intergenic region. All cloning, including introduction of temperature-sensitive mutations, was performed via standard and overlapping PCRs with CloneAmp™ HiFi PCR Premix (Takara Bio, cat # 639298, San Jose, CA), with subsequent insertion into the construct at unique restriction sites by In-Fusion ligation-independent cloning (Takara Bio, San Jose, CA). All cloning was performed with Stbl2 *E. coli* (Invitrogen) with growth at 30 °C. All guides were designed as described using the CRISPR design tool (crispr.mit.edu)^68^ in coordination with our in-house RNA folding prediction pipeline. The T7-driven helper plasmids encoding SeV-N, SeV-P, and SeV-L were the kind gift of Nancy McQueen. Rescue of Sendai virus was performed as described previously^36,65,69^ by transfecting with 4 μg T7-driven antigenome, 1.44 μg T7-N, 0.77 μg T7-P, 0.07 μg T7-L, and 4 μg codon-optimized T7 polymerase, using Lipofectamine LTX (Invitrogen) according to manufacturer’s protocol. After rescue and amplification, supernatant was clarified the purified by ultracentrifugation in a discontinuous 20% to 65% sucrose gradient allowing the interface to be collected.

### Flow cytometry

For CCR5 staining, cells were lifted and blocked in phosphate-buffered saline with 2% FBS. Alexa 647-conjugated rat anti-human CCR5 (cat# 313712, BioLegend, San Diego, CA) was added at 1:100 for 30 minutes at 4 °C before washing and resuspension in 2% paraformaldehyde. For p24 staining (RD1-conjugated mouse anti-p24 clone KC57, cat# 6604667, 1:100 dilution, Beckman Coulter, Brea, CA), cells were fixed and permeabilized using the Cytofix/Cytoperm kit (BD Biosciences, San Jose, CA) before blocking. Flow cytometry was performed on a BD LSR II at the Flow Cytometry Core at the Icahn School of Medicine at Mount Sinai.

### Characterization of editing efficiency

Genomic DNA was extracted using the PureLink Genomic DNA Mini Kit (Invitrogen). Specific genomic loci were amplified using Velocity DNA Polymerase (Bioline). Off-target loci represent the top predicted off-target sites in the CRISPR Design Tool (crispr.mit.edu)^68^. PCR products were gel-extracted (NucleoSpin Gel and PCR Clean-up kit, Clontech) and sent for Sanger sequencing. Sequencing results could then be uploaded to the Synthego ICE Analysis tool (v3) allowing for inference of the percent indels in the sample. For deep sequencing, the gel-extracted products were pooled and prepared for sequencing via paired-end 2 × 150 bp iSeq (Illumina, San Diego, CA) sequencing in-house.

### Human CD34+ HSPCs from mobilized peripheral blood and fetal liver

Fetal livers were obtained from the UCLA Center for AIDS Research (CFAR) Gene and Cellular Therapy Core. The UCLA institutional review board has determined that these tissues are not human subjects and do not require an institutional review board review, because fetal tissues were obtained without patient-identifying information from deceased fetuses. Written informed consent was obtained from patients for the use of tissues in research purposes. CD34^+^ HSPCs were isolated from fetal livers using anti-CD34^+^ magnetic bead-conjugated monoclonal antibodies (Miltenyi Biotec) and cryopreserved in Bambanker (Wako Chemical USA).

### CD14+ infection methods

Leukopaks were purchased from the New York Blood Bank and CD14+ monocytes were isolated with the EasySep Human CD14 positive selection kit (StemCell #17858). CD14+ monocytes were mock-infected or infected with ts SeV-Cas9-US11 or ts SeV-Cas9-CCR5 virus at an MOI of 10. Cells were incubated with virus for 1 hour at 37°C in a microfuge tube rotating rack, to ensure even distribution of cells and virus. After the inoculation, cells were briefly pelleted and resuspended in R10 medium (RPMI supplemented with FBS, HEPES, L-glutamine, and penicillin-streptomycin) with 50 ng/mL of GMSF (Sigma Aldrich G5035) and were seeded into a 24-well plate at a density of 1E+06 cells/mL and incubated at 34°C, with 6 wells per sample. Media was replaced with fresh GM-CSF every 3 days to differentiate the CD14+ monocytes into macrophages. The cells were shifted to 37°C at 3 days post infection.

At 7 days post temperature shift, the monocyte-derived macrophages (MDMs) were imaged using the Celigo Imaging cytometer (Nexcelom) to verify SeV-Cas9 infection by GFP+ expression. Genomic DNA from 2 wells per sample was isolated using the PureLink Genomic DNA mini kit (ThermoFisher #K182001). The remaining MDMs were infected with HIV strain JFRL-mCherry at 10,000 pg of P24 per million cells by spinoculation at 1200 rpm for 2 hours at room temperature. Cells were rinsed 1 time with PBS to remove inoculum, R10 media with GM- CSF was added, and the MDMs were incubated at 37°C. At 1, 4, and 8 dpi, 200 µL of cell supernatant was removed and frozen for P24 analysis and the media was replaced in each well to maintain a total volume of 500 µL per well. At 8 dpi with HIV, the MDMs were imaged again with the Celigo Imaging cytometer. Genomic DNA was isolated from each sample. *CCR5* editing efficiency was determined by PCR amplification of the *CCR5* locus, sanger sequencing, and Synthego ICE analysis. P24 levels in the supernatant of HIV-infected MDMs was determined by ELISA (Xpress Bio HIV-1 p24 ELISA Assay).

### CD14+ infection and HIV challenge

Leukopaks were purchased from the New York Blood Bank and CD14+ monocytes were isolated with the EasySep Human CD14 positive selection kit (StemCell #17858). CD14+ monocytes were mock-infected or infected with ts SeV-Cas9-US11 or ts SeV-Cas9-CCR5 virus at an MOI of 10. Cells were incubated with virus for 1 hour at 37°C in a microfuge tube rotating rack, to ensure even distribution of cells and virus. After the inoculation, cells were briefly pelleted and resuspended in R10 medium (RPMI supplemented with FBS, HEPES, L-glutamine, and penicillin-streptomycin) with 50 ng/mL of GMSF (Sigma Aldrich G5035) and were seeded into a 24-well plate at a density of 1E+06 cells/mL and incubated at 34°C, with 6 wells per sample. Media was replaced with fresh GM-CSF every 3 days to differentiate the CD14+ monocytes into macrophages. The cells were shifted to 37°C at 3 days post infection.

At 7 days post temperature shift, the monocyte-derived macrophages (MDMs) were imaged using the Celigo Imaging cytometer (Nexcelom) to verify ts SeV-Cas9 infection by GFP+ expression. Genomic DNA from 2 wells per sample was isolated using the PureLink Genomic DNA mini kit (ThermoFisher #K182001). The remaining MDMs were infected with HIV strain JRFL-mCherry^70–72^ at 10,000 pg of P24 per million cells by spinoculation at 1200 rpm for 2 hours at room temperature. Cells were rinsed 1 time with PBS to remove inoculum, R10 media with GM-CSF was added, and the MDMs were incubated at 37°C. At 1, 4, and 8 dpi, 200 µL of cell supernatant was removed and frozen for P24 analysis and the media was replaced in each well to maintain a total volume of 500 µL per well. At 8 dpi with HIV, the MDMs were imaged again with the Celigo Imaging cytometer. Genomic DNA was isolated from each sample. CCR5 editing efficiency was determined by PCR amplification of the CCR5 locus, sanger sequencing, and Synthego ICE analysis. P24 levels in the supernatant of HIV-infected MDMs was determined by ELISA (Xpress Bio HIV-1 p24 ELISA Assay).

## ACKNOWLEDGMENTS

The authors acknowledge the following funding: CS was supported by Viral-Host Pathogenesis Training Grant T32 AI07647; BL acknowledges flexible funding support from NIH grants R01 AI123449, R21 AI1498033, and the Department of Microbiology and the Ward-Coleman estate for endowing the Ward-Coleman Chairs at the ISMMS. HIV JR-FL was obtained through the NIH AIDS Reagent Program, Division of AIDS, NIAID, NIH: HIV-1 JR-FL Virus from Dr. Irvin Chen. .

---

Supplemental Figure 1. The editing efficiency of ts-SeV-Cas9 mCherry in CD34+ HSCs. Indel frequency calculated as described in Figure 3B.

Supplemental Figure 2. The effect on hematopoietic differentiation of CD34+ HSPCs following infection by SeV Cas9. Raw colony counts, performed as described in Figure 5C.

## Declaration of Generative AI and AI-assisted technologies in the writing process

During the preparation of this work the authors used OpenAI’s ChatGPT in order to proof work for clarity of writing. After using this tool/service, the authors reviewed and edited the content as needed and take full responsibility for the content of the publication.

## Notes

### Competing Interest Statement

The authors have declared no competing interest.

