## Supplementary Figures for "A temperature-sensitive and interferon-silent Sendai virus vector for CRISPR-Cas9 delivery and gene editing in primary human cells"

### Slide 1
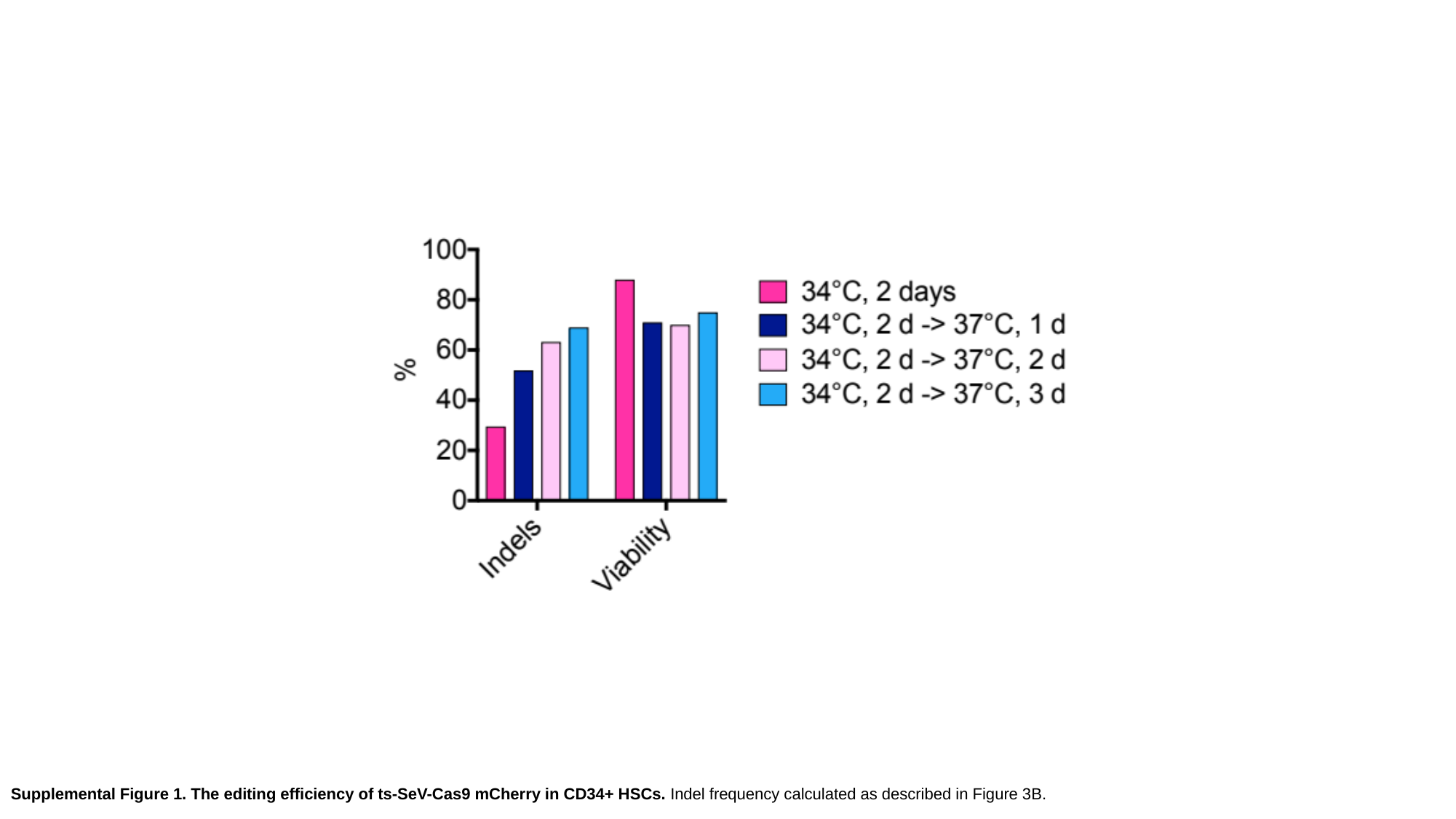

Supplemental Figure 1. The editing efficiency of ts-SeV-Cas9 mCherry in CD34+ HSCs. Indel frequency calculated as described in Figure 3B.

### Slide 2
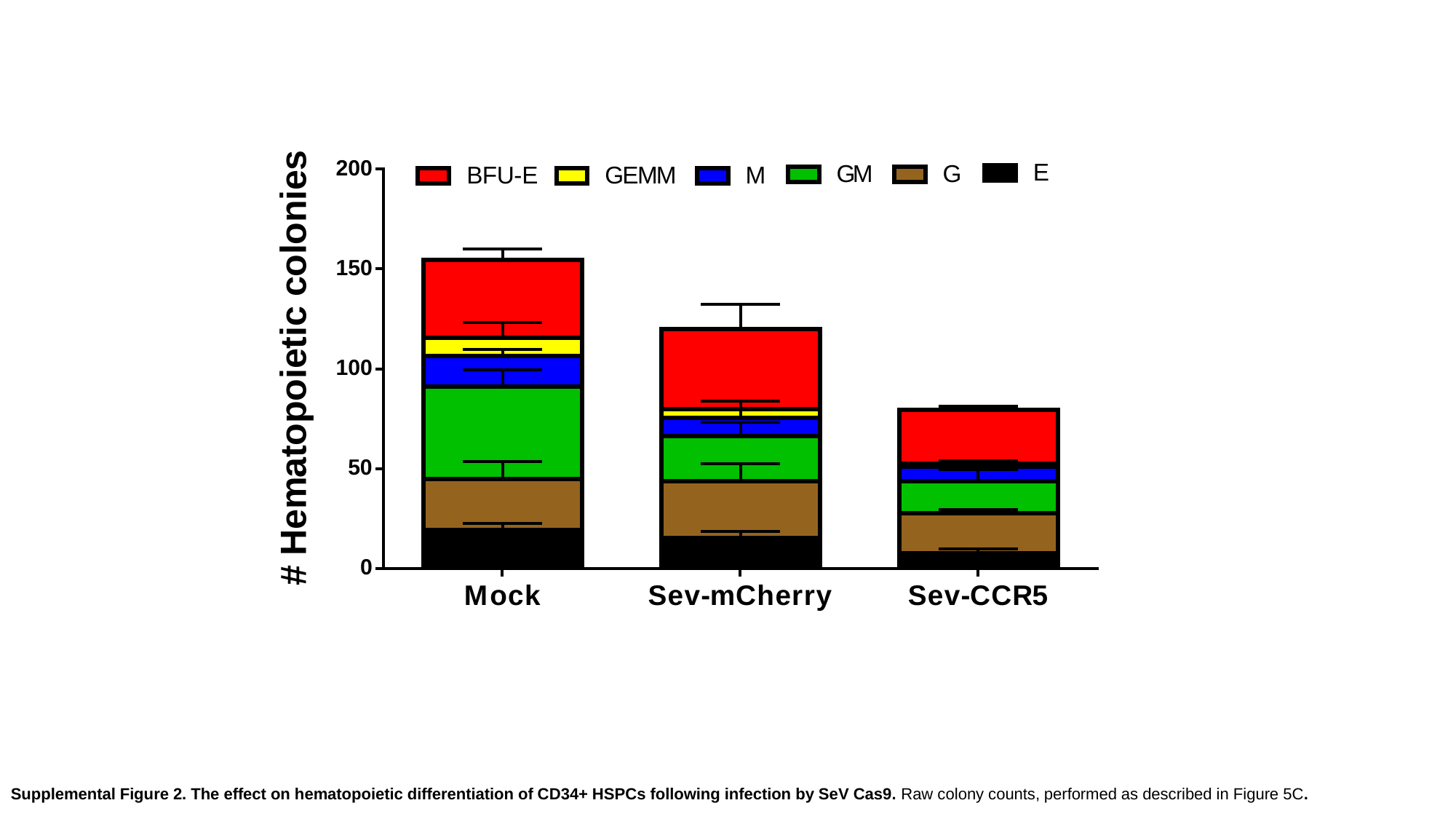

Supplemental Figure 2. The effect on hematopoietic differentiation of CD34+ HSPCs following infection by SeV Cas9. Raw colony counts, performed as described in Figure 5C.
